# The unfolded protein response of the endoplasmic reticulum supports mitochondrial biogenesis by buffering non-imported proteins

**DOI:** 10.1101/2021.05.19.444788

**Authors:** Katharina Knöringer, Carina Groh, Lena Krämer, Kevin C. Stein, Katja G. Hansen, Johannes M. Herrmann, Judith Frydman, Felix Boos

## Abstract

Almost all mitochondrial proteins are synthesized in the cytosol and subsequently targeted to mitochondria. The accumulation of non-imported precursor proteins occurring upon mitochondrial dysfunction can challenge cellular protein homeostasis. Here we show that blocking protein translocation into mitochondria results in the accumulation of mitochondrial membrane proteins at the endoplasmic reticulum, thereby triggering the unfolded protein response (UPR^ER^). Moreover, we find that mitochondrial membrane proteins are also routed to the ER under physiological conditions. The levels of ER-resident mitochondrial precursors is enhanced by import defects as well as metabolic stimuli that increase the expression of mitochondrial proteins. Under such conditions, the UPR^ER^ is crucial to maintain protein homeostasis and cellular fitness. We propose the ER serves as a physiological buffer zone for those mitochondrial precursors that can’t be immediately imported into mitochondria while engaging the UPR^ER^ to adjust the ER proteostasis capacity to the extent of precursor accumulation.

## Introduction

The ability of cells to maintain protein homeostasis (proteostasis) is crucial for organismal health. Imbalances in protein synthesis, targeting, folding and degradation are associated with numerous diseases and are also hallmarks of aging ^1-5^. Cells constantly monitor their proteome to quickly sense proteotoxic perturbations and launch stress-reactive programs to restore homeostasis. Of particular importance are the compartment-specific stress responses to misfolded proteins of the cytosol and nucleus (heat shock response) as well as of the endoplasmic reticulum (unfolded protein response of the ER, UPR^ER^) and mitochondria (UPR^mt^). Via the activation of dedicated transcription factors, these pathways elevate the levels of chaperones, proteases and other quality control factors in the compartment where protein misfolding is sensed ^6-9^.

When misfolded proteins accumulate in the ER, the ER membrane kinase Ire1 dimerizes, autophosphorylates and then splices the mRNA of *XBP1* (*HAC1* in yeast). This enables its efficient translation, giving rise to a potent transcription factor that induces the UPR^ER 10^. Besides increasing the expression of ER chaperones and other biogenesis factors, the UPR^ER^ can considerably expand the ER of a cell. In yeast, the Ire1-Hac1 pathway is the only dedicated regulator of the UPR^ER^, while mammalian cells have two additional branches of the UPR^ER^ that control transcription, translation and eventually apoptosis via PERK and ATF6 ^11^.

Cellular organelles have clearly distinct organizations and functions, yet they are no independent entities; instead, they form tight physical contacts ^12-14^ and functionally cooperate in the synthesis of proteins, lipids and metabolites ^15,16^. Hence, they mutually influence and rely on the homeostasis of one another. In many protein folding diseases, defects in proteostasis are observed in multiple organelles at the same time, even though the primary perturbation occurs in most cases in only one compartment ^17,18^. As a consequence, the different stress response programs need to act in concert ^19-23^. For instance, perturbations of mitochondrial proteostasis often compromise mitochondrial protein import so that non-imported precursor proteins accumulate in the cytosol ^19,24-26^. Consequently, mitochondrial dysfunction not only activates mitochondrial quality control pathways, but also the expression of cytosolic chaperones and the ubiquitin-proteasome system, which mitigate the deleterious effects of mistargeted precursors ^27-29^. In addition, the synthesis of many mitochondrial proteins is muted by transcriptional repression as well as global translation attenuation to further reduce the burden on cytosolic proteostasis ^28,30,31^.

While numerous pathways of cross-compartment communication under proteotoxic stress have been identified, our understanding of the connections between organellar stress response programs is still incomplete. Here we show that defective mitochondrial protein import not only activates mitochondrial and cytosolic stress responses, but also triggers the unfolded protein response of the ER. This is at least in part attributable to the targeting of mitochondrial membrane proteins to the ER. The UPR^ER^ is functionally relevant both under conditions of compromised protein import, and conditions that induce mitochondrial biogenesis such as metabolic adaptations. Thus, the UPR^ER^ supports mitochondrial biogenesis by buffering the adverse consequences of elevated levels of non-imported mitochondrial precursor proteins.

## Results

### The unfolded protein response of the ER is triggered by long-lasting mitoprotein-induced stress

Cellular adaptations to imbalances in mitochondrial proteostasis have been studied using mutants of protein import components ^27^, chaperones ^29^, folding-incompetent mitochondrial proteins ^31,32^, or defects in the respiratory chain ^33,34^. Many of these perturbations converge on the impairment of mitochondrial protein import. Model systems in which protein import can be acutely blocked have proven particularly useful to decipher the mechanistic details of responses to such mitoprotein-induced stress. A way of achieving this is the overexpression of mitochondrial precursor proteins that are intrinsically prone to premature folding and stalling inside the narrow mitochondrial translocases ^35^. For instance, the well-characterized ‘clogger’ protein *b*_2_-DHFR can be used for this purpose ^28,36^. This fusion protein consists of the N-terminal 167 amino acids of cytochrome *b*_2_ (including its mitochondrial targeting signal) and the rapidly and tightly folding dihydrofolate reductase DHFR (Fig. 1A) ^37-39^. Expression of the clogger results in accumulation of non-imported precursor proteins (Fig. 1B) and impairs cell growth (Fig. 1C). In baker’s yeast, the expression of *b*_2_-DHFR can be tightly controlled by using a *GAL* promoter that can be switched on by the addition of galactose to the lactate-based growth media. This allows for a tight temporal resolution and the discrimination between short-term and long-term responses to an acute and specific blockade of protein import.

**Figure 1.**
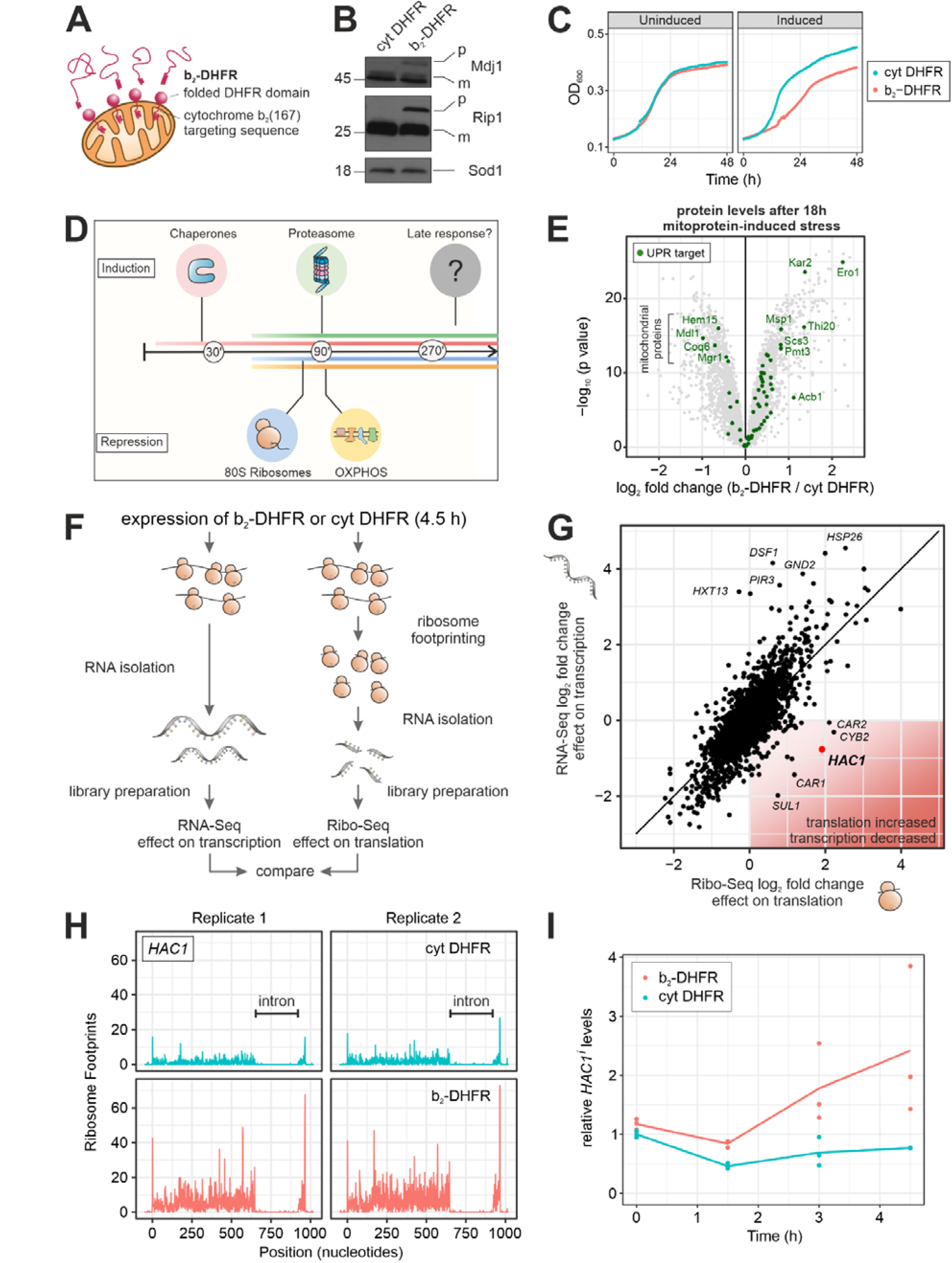
Mitoprotein-induced stress triggers the UPR^ER^. **A**, Fusion of DHFR to the N-terminus of Cytochrome *b*_*2*_ generates a mitochondrial ‘clogger’ that jams the protein import machinery. **B**, The mitochondrial clogger *b*_*2*_-DHFR or cytosolic DHFR were expressed for 4.5 h. The precursor form of the mitochondrial proteins Mdj1 and Rip1 were detected by Western Blotting. **C**, Expression of *b*_*2*_-DHFR leads to attenuated growth. **D**, The mitoprotein-induced stress response encompasses an early transcriptional induction of chaperones and the proteasome and a downregulation of cytosolic ribosomes and OXPHOS components. **E**, Protein levels in clogger-expressing versus control cells after 18 h of induction were measured by quantitative mass spectrometry ^28^. Highlighted are proteins which are reported targets of the UPR^ER 21^. Data from *n*=3 independent biological replicates are shown. **F, G**, The cellular transcriptome and translatome after 4.5 h of clogger induction were measured by RNA-Seq (*n*=4) ^28^ and ribosome profiling (*n*=3), respectively. Shown are log_2_ fold changes of *b*_*2*_-DHFR versus cytosolic DHFR. *HAC1* transcripts are slightly reduced, but its translation is upregulated. **H**, Ribosome footprints along the *HAC1* gene from cells expressing *b*_*2*_-DHFR or cytosolic DHFR for 4.5 h are shown. **I**, Levels of spliced *HAC1* mRNA in cells expressing *b*_*2*_-DHFR or cytosolic DHFR were measured by RT-qPCR over time (*n*=3).

We previously characterized the immediate reactions of the cellular transcriptome to mitoprotein-induced stress ^28^. An induction of many chaperones and the proteasome and a repression of OXPHOS components and ribosomes all took place within 1.5 h of clogger expression, some of them even markedly earlier (Fig. 1D). However, many cellular adaptations change when acute stress persists and becomes long-lasting ^40-42^. We therefore asked whether cells undergo additional adaptations when exposed to long-term mitoprotein-induced stress. To this end, we reanalyzed our previously collected data to examine changes in the cellular proteome after up to 18 h of clogger expression ^28^. We queried for changes in the proteome that were evident at time points no earlier than 4.5 h, which corresponds to approximately one cell doubling in respiratory medium. Interestingly, this criterion identified a group of proteins that are associated with the unfolded protein response of the ER (Fig. 1E and S1A) ^21^. Some individual targets of the UPR^ER^, such as e.g. Ero1 and Kar2, were induced at earlier time points, presumably due to their responsiveness to the transcription factors Hsf1 and/or Rpn4 that form the first line of defense against mitoprotein-induced stress. Moreover, a small number of UPR targets were decreased over time. These proteins (Hem15, Mdl1, Coq6, Mgr1) almost exclusively localize to mitochondria and their levels are likely affected by the import block or the clogger-induced downregulation of mitochondrial components. However, most UPR targets showed a consistent upregulation that was observed 9 h after clogger induction, and even more so after 18 h (Fig. 1E and S1A).

In yeast, the UPR^ER^ is activated by splicing of an intron from the *HAC1* mRNA in the cytosol through the ER-resident kinase Ire1. Only the spliced isoform of the mRNA (called *HAC1*^*i*^) can be translated and gives rise to a transcription factor ^10^. To test whether *HAC1* was indeed spliced and translated under mitoprotein-induced stress, we analyzed clogger-expressing cells by ribosome profiling. Here, ribosome footprints from cells expressing *b*_*2*_-DHFR or cytosolic DHFR were sequenced 4.5 h after induction, and the changes in the translatome were compared to the changes in the transcriptome (Fig. 1F). For the large majority of all genes, transcriptional and translational changes correlated tightly. For *HAC1* however, we observed a slight reduction of mRNA levels, while we found four times more ribosome footprints on *HAC1* mRNA in clogger-expressing than in control cells (Fig 1G). In fact, *HAC1* was one of the most prominent outliers in this comparison, ranking as the gene with the second-highest gain in translational efficiency when mitochondrial import was blocked (Fig. S1B). The increase in ribosome occupancy was restricted to the exon region of the mRNA, while the intron region of *HAC1* was free of ribosome densities in both conditions (Fig 1H).

We next sought to more precisely determine the timing of the UPR^ER^ activation. To this end, we set up an RT-qPCR assay which quantifies the spliced isoform of *HAC1*^*i*^ by using a primer-probe combination which specifically recognizes the exon-exon junction of *HAC1*^*i*^ (Fig. S1C, D). We induced *b*_*2*_-DHFR by addition of 0.5% galactose to cultures that were previously grown in lactate medium and followed *HAC1* splicing over time. The earliest time point at which we could detect a considerable difference between clogger-expressing and control cells was 3 h (Fig. 1I). As a certain delay between the onset of *HAC1* splicing and downstream changes in protein levels of UPR^ER^ targets is expected, this is consistent with our earlier observation that UPR^ER^ induction is a rather late event in mitoprotein-induced stress signaling.

We conclude that under long-term impairment of mitochondrial protein import, cells induce the UPR^ER^ via the canonical Ire1-Hac1 pathway.

### UPR^ER^ induction is required for cellular fitness under mitoprotein-induced stress

We asked whether UPR^ER^ induction is functionally relevant under sustained mitoprotein-induced stress, given that its magnitude is rather mild when compared to harsh ER insults such as treatment with tunicamycin (*cf*. Fig. S1C). To this end, we compared the fitness of UPR^ER^-deficient cells with that of wild type cells when mitochondrial import was blocked. Indeed, when either *HAC1* or *IRE1* were deleted, cells exhibited synthetic growth defects upon clogger expression, both in liquid medium and on plates (Fig. 2A, B).

**Figure 2.**
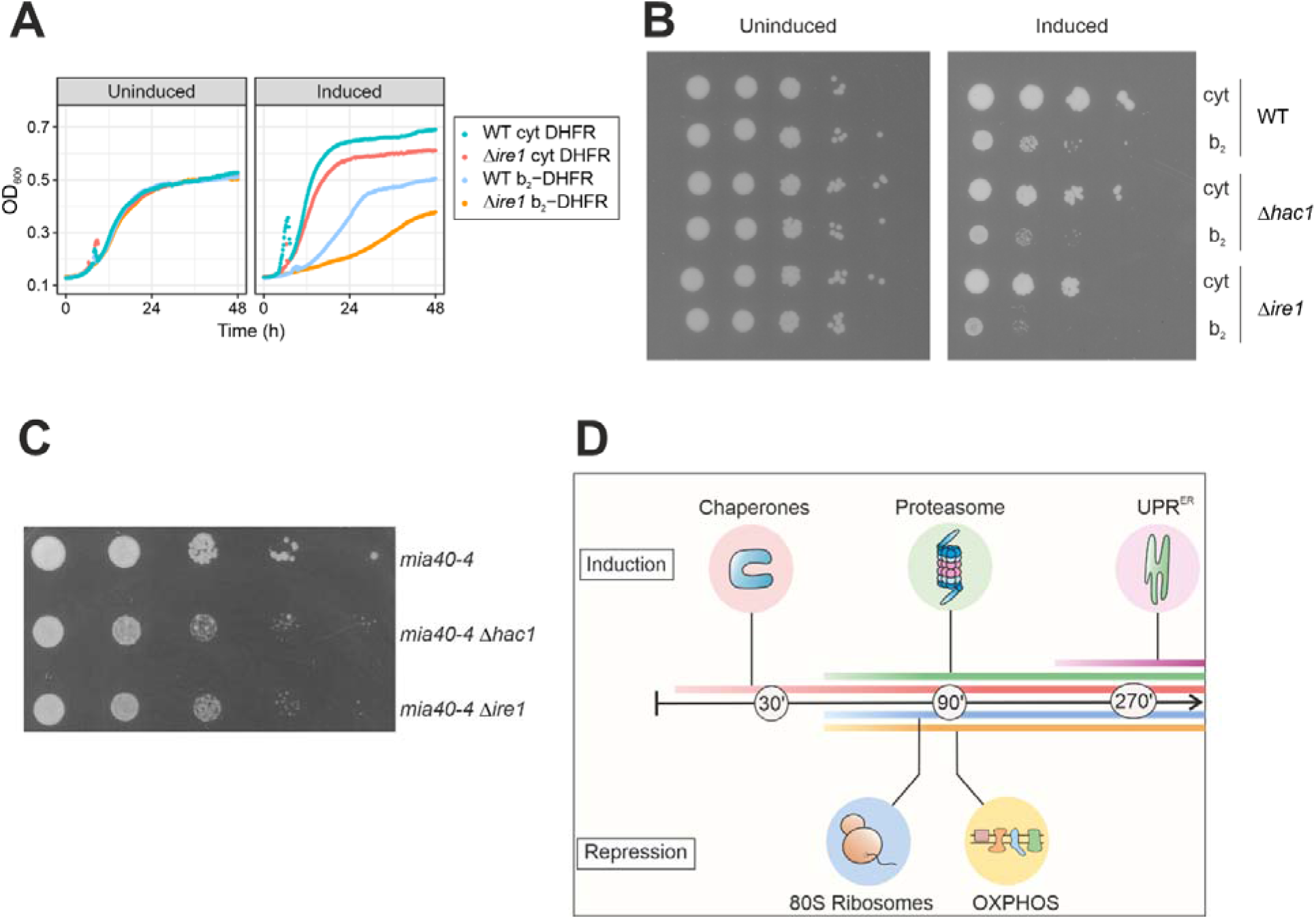
The UPR^ER^ is required for cellular fitness under mitoprotein-induced stress conditions. **A**, Wild type and Δ*ire1* cells were grown under non-inducing (left) or inducing (right) conditions, expressing either *b*_*2*_-DHFR or cytosolic DHFR. Δ*ire1* cells are more susceptible to mitoprotein-induced stress. **B**, Tenfold serial dilutions of wild type, Δ*ire1* and Δ*hac1* cultures were dropped on lactate plates with (‘induced) or without (‘uninduced’) 0.5% galactose. The UPR^ER^-deficient mutants show synthetic growth defects with expression of *b*_*2*_-DHFR. **C**, *HAC1* and *IRE1* were deleted in temperature-sensitive *mia40-4* mutants. Cells were grown in glucose medium and serial dilutions were spotted on glucose plates and incubated at the semi-permissive temperature of 30°C. Loss of the UPR^ER^ results in synthetic growth defects. **D**, Early cytonuclear adaptations to mitoprotein-induced stress are accompanied by the induction of the UPR^ER^ as a second line of defense.

We examined the relevance of UPR^ER^ signaling when mitochondrial import is impaired by an approach orthogonal to clogging the translocases. To this end, we deleted *HAC1* or *IRE1* in a strain that carries a temperature-sensitive mutation in the essential import component Mia40. Mia40 is responsible for the import and oxidative folding of cysteine-containing mitochondrial intermembrane space proteins ^43,44^. The import defects in the *mia40-4* mutant were shown to trigger cytosolic adaptations (unfolded protein response activated by mistargeting of proteins, UPR^am^) similar to those elicited by the clogger ^27^. Indeed, *mia40-4* cells grew worse at semi-permissive growth conditions when *IRE1* or *HAC1* were deleted, demonstrating that UPR^ER^ signaling is relevant when protein import into the IMS is perturbed (Fig. 2C).

In conclusion, defects in mitochondrial protein import trigger the UPR^ER^, which is required for cellular fitness under such conditions (Fig. 2D).

### Mitochondrial membrane proteins accumulate at the ER when mitochondrial protein import is impaired

What could be the cause for UPR^ER^ activation in situations when mitochondrial import is impaired? Blocking import should elevate levels of mitochondrial precursor proteins in the cytosol. Therefore, we reasoned that a portion of these non-imported precursors, perhaps comprising membrane proteins, may be targeted to the ER, where they would accumulate and engage folding and protein quality control systems, thus triggering UPR^ER^ activation.

To test this hypothesis, we labelled the mitochondrial inner membrane protein Oxa1 with ymNeonGreen, and coexpressed it with Sec63-ymScarletI as an ER marker, followed by analysis of their subcellular distribution by fluorescence microscopy. When we expressed cytosolic DHFR, the green Oxa1 signal and the red Sec63 signal partitioned into separate structures with no considerable colocalization. In contrast, when *b*_*2*_-DHFR was expressed for 4.5 h, we found that a fraction of Oxa1-ymNeonGreen colocalized with Sec63-ymScarletI in the typical ring-shaped structures of the perinuclear and peripheral ER (Fig. 3A). This ER co-localization was observed in around 30% of the clogger-expressing cells, but only in around 1% of control cells (Fig. 3B).

**Figure 3.**
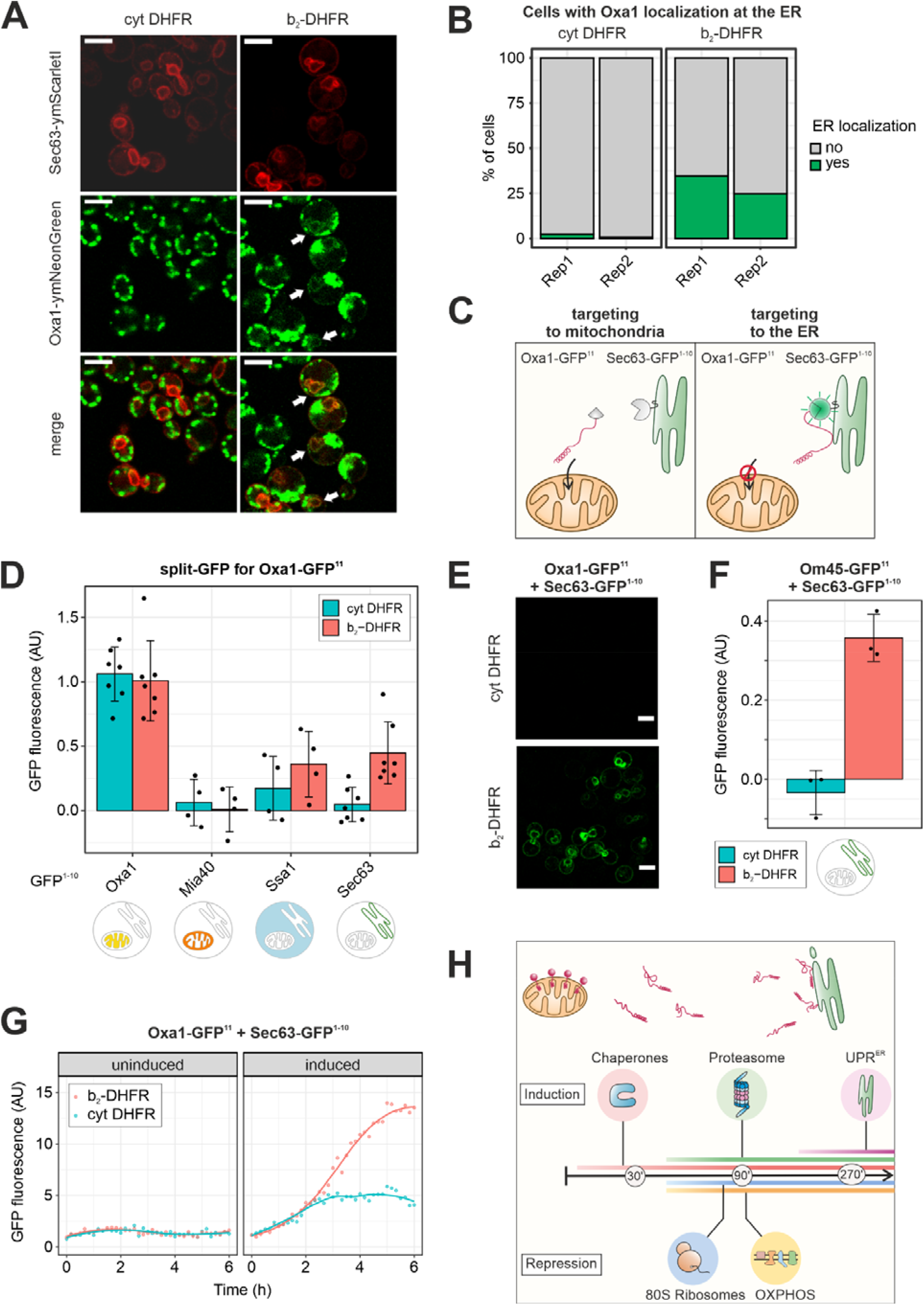
Non-imported mitochondrial membrane proteins localize to the ER. **A**, The mitochondrial inner membrane protein Oxa1 was genomically tagged with ymNeonGreen, the ER marker Sec63 was tagged with ymScarletI. Confocal fluorescence microscopy was performed after 4.5 h of expression of either *b*_*2*_-DHFR or cytosolic DHFR. When the clogger was induced, a fraction of Oxa1-ymNeonGreen colocalized with Sec63-ymScarletI. Scale bar, 5 µm. **B**, Quantification of the number of cells from A in which ER localization of Oxa1-ymNeonGreen was observed. **C**, Schematic depiction of the split-GFP strategy to measure ER localization of mitochondrial proteins. **D**, The GFP^11^ fragment was fused to Oxa1 and the GFP^1-10^ fragment was fused to Oxa1, Mia40, Ssa1 or Sec63. *b*_*2*_-DHFR or cytosolic DHFR were induced for 4.5 h and fluorescence was measured in a platereader. Mean values and standard deviations are shown for *n*=7 (Oxa1-GFP^1-10^, Sec63-GFP^1-10^) or *n*=4 (Mia40-GFP^1-10^, Ssa1-GFP^1-10^) independent biological replicates. **E**, Fluorescence microscopy of cells expressing Oxa1-GFP^11^ and Sec63-GFP^1-10^ and either *b*_*2*_-DHFR or cytosolic DHFR after 4.5 h of induction. Scale bar, 5 µm. **F**, The GFP^11^ fragment was fused to Om45 and the GFP^1-10^ fragment to Sec63. Clogger expression for 4.5 h evoked an increase in fluorescence (mean values and standard deviations for *n*=3 independent biological replicates). **G**, Cells expressing Oxa1-GFP^11^ and Sec63-GFP^1-10^ were cultured in lactate medium before either *b*_*2*_-DHFR or cytosolic DHFR were induced by addition of 0.5 % galactose. Fluorescence was monitored in a Clariostar plate reader every 10 min for *n*=6 biological replicates. Constitutively expressed ymScarletI was used to normalize for growth and overall translation rates. After around 3 h of induction, elevated split-GFP signals in clogger-expressing cells indicated accumulation of Oxa1 at the ER. **H**, Model for the connection between mitochondrial import block and UPR^ER^ induction. Clogging the mitochondrial translocases leads to accumulation of precursor proteins in the cytosol as well as at the ER surface, which triggers the ER stress response.

Fusions with fluorescent proteins can interfere with the function, localization, folding and stability of proteins ^45-47^. In particular, a large, stably folding C-terminal moiety might generate a mitochondrial clogger, as exemplified by *b*_*2*_-DHFR itself, and interfere with import and localization. We therefore sought to verify our results by a method that avoids the fusion of large protein domains to mitochondrial precursors and also minimizes the need for manual categorization of microscopic images with high mitochondrial background signal. To this end, we adapted a split-GFP method specifically designed to assess protein localization *in vivo* ^48^. Superfolder GFP is split into two parts, GFP^1-10^ and GFP^11^, which only emit fluorescence when co-localized to the same compartment (Fig. 3C). The GFP fragments do not alter the folding behavior of the fusion proteins and their affinity is high enough to promote self-association without the need for a direct protein-protein interaction of the fusion partners ^49-51^. The GFP^11^ tag consists of only 17 amino acid residues and is therefore unlikely to affect translocation across the mitochondrial membranes.

To first verify that the split-GFP assay captures the subcellular localization of mitochondrial proteins, we fused the GFP^11^ fragment to the C-terminus of Oxa1 (inner membrane, C-terminus at matrix side), Mia40, Dld1 (inner membrane, C-terminus at IMS side) and Om45 (outer membrane, C-terminus at IMS side) and the GFP^1-10^ fragment to Oxa1 (IM, matrix side), Mia40 (IM, IMS side), Sec63 (ER membrane, cytosolic side) and Ssa1 (cytosol) (Fig. S2A) ^52-58^. In the absence of any stress, the by far strongest fluorescence signal was detected for the combinations that recapitulate the known localization and topology for all proteins tested (Oxa1-GFP^11^ / Oxa1-GFP^1-10^, Mia40-GFP^11^ / Mia40-GFP^1-10^, Dld1-GFP^11^ / Mia40-GFP^1-10^, Om45-GFP^11^ / Mia40-GFP^1-10^), while all other combinations resulted in much lower fluorescent signals (Fig. 3D and S2B-D, control condition in blue). This showed that the approach can measure protein localization with sub-organellar resolution.

We next expressed either *b*_*2*_-DHFR or cytosolic DHFR for 4.5 h in strains carrying the split-GFP reporters. *b*_*2*_-DHFR expression evoked a marked increase in signal for Oxa1-GFP^11^ with the Sec63-GFP^1-10^ and the Ssa1-GFP^1-10^ reporters, while with cytosolic DHFR, only very little signal was detected (Fig. 3D). This points towards relocation of a fraction of newly synthesized Oxa1-GFP^11^ to the ER surface or, potentially, the cytosol. We used fluorescence microscopy to confirm that the fluorescence we measured in a plate reader setup indeed originated from ER-localized GFP complementation (Fig. 3E and S2E-F). For the mitochondrial outer membrane protein Om45-GFP^11^, we found a similar redistribution to the ER under import stress (Fig. 3F and S2B), while neither Mia40-GFP^11^ nor Dld1-GFP^11^ showed detectable ER localization (Fig. S2C, D). Obviously, some but not all mitochondrial membrane proteins are routed to the ER when their entry into mitochondria is delayed.

How is the timing of precursor localization to the ER after mitochondrial import is blocked? To assess this question, we grew cells expressing Oxa1-GFP^11^ and Sec63-GFP^1-10^ in a plate reader, induced *b*_*2*_-DHFR or cytosolic DHFR by addition of galactose and monitored split-GFP fluorescence over time in living cells. Constitutively expressed ymScarletI was used to normalize for differences in cell growth and translation rates. Clogger-expressing cells showed elevated split-GFP signals from around 3 h after induction (Fig. 3G). Strikingly, the induction of the UPR^ER^ and the detection of Oxa1 at the ER perfectly coincided in time (*cf*. Fig. 1I).

In conclusion, when mitochondrial import is blocked, some mitochondrial preproteins accumulate at the ER membrane which likely evokes the UPR^ER^ (Fig. 3H).

### The UPR^ER^ maintains cellular fitness during adaptation of mitochondrial biogenesis

Is the UPR^ER^ only a stress-reactive system that comes into play when mitochondrial import is defective, or is it of more general relevance for mitochondrial biogenesis? Accurate protein sorting is a challenging task for cells, and the ER might constantly encounter a certain load of mitochondrial precursor proteins. To check whether there is evidence for mitochondrial proteins routed to the ER in the absence of stress, we reanalyzed several high-resolution datasets on protein targeting. Proximity labeling of ribosomes close to the ER or the mitochondrial outer membrane and subsequent ribosome profiling determined the ‘local translatome’ at the ER and mitochondrial surface in yeast ^59,60^. Interestingly, while most mitochondrial proteins were enriched in the vicinity of mitochondria, a subset of mitochondrial proteins was found to be translated close to the ER, notably including Oxa1 (Fig. 4A). Also in human cells, mRNAs of some mitochondrial proteins were found at the ER surface (Fig. S3A) ^61^. Finally, we reanalyzed datasets from studies that determined which nascent chains interact with the signal recognition particle (SRP) in yeast by pulldown of SRP and subsequent sequencing of the bound transcripts ^62,63^. SRP is a major targeting factor for secretory proteins that carry a signal sequence or transmembrane domains ^64-67^. While secretory proteins were clearly the most enriched among the SRP substrates, a subset of mitochondrial encoding ribosome-nascent chains were also bound by SRP to a lesser extent, but significantly above what was found for cytosolic proteins (Fig. S3B). Both in yeast and in human cells, ER-localized mitochondrial transcripts include, but are not limited to proteins with known dual localization to mitochondria and ER. Apparently, some mitochondrial precursors have a tendency to be targeted to the ER even in the absence of stress, possibly mediated by ‘low priority’ SRP-binding to at least some of these precursors.

**Figure 4.**
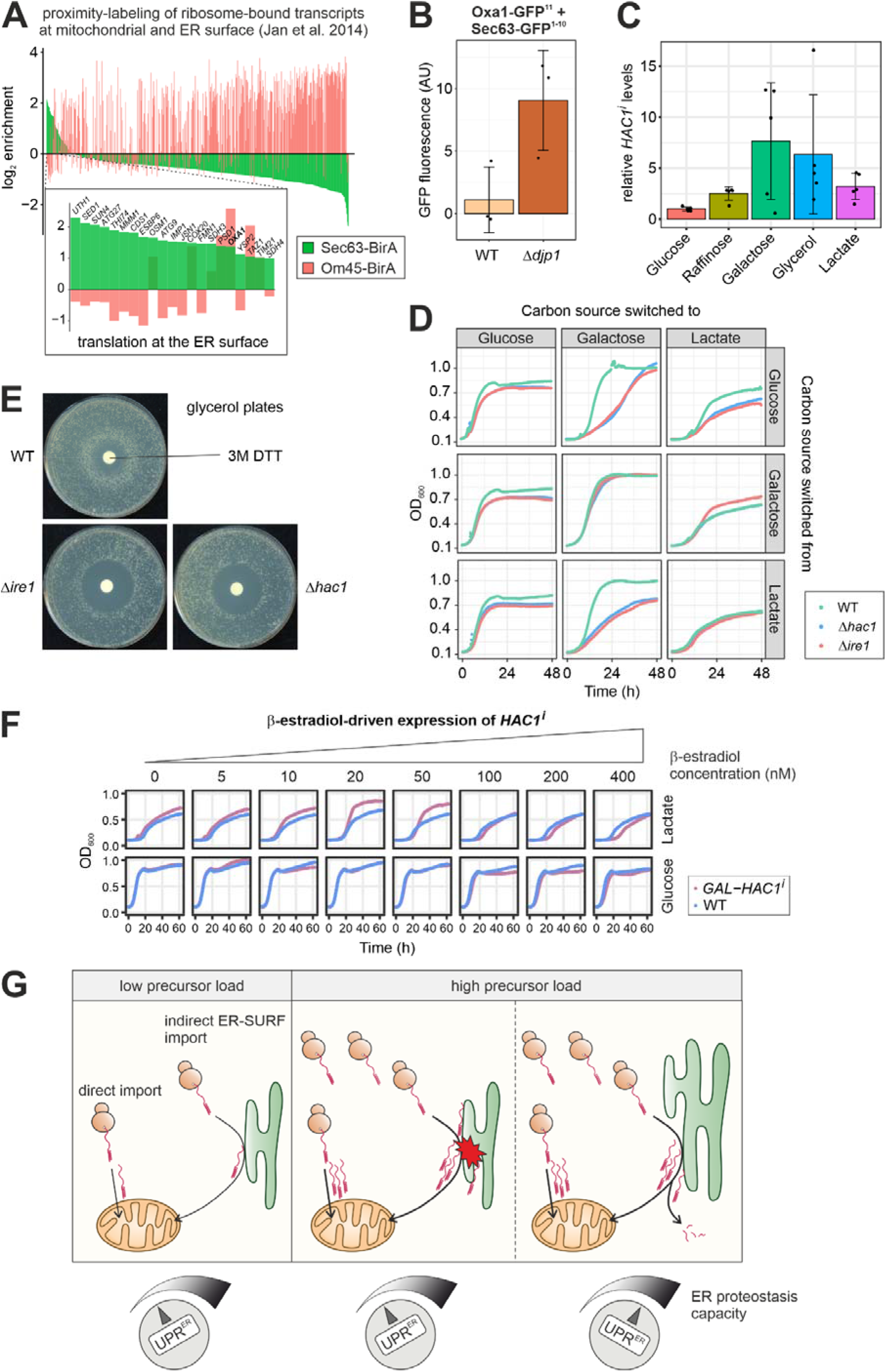
The UPR^ER^ maintains cellular fitness during changes in mitochondrial biogenesis. **A**, Data from Jan et al. (2014)^59^ on the localized translation near the mitochondrial and the ER surface. For all mitochondrial proteins in the dataset, the log_2_ enrichment of ribosome-nascent chain complexes at the ER membrane (Sec63-BirA) and the mitochondrial outer membrane (Om45-BirA) over the total ribosomes are shown. While most translated mRNAs localize to the mitochondrial membrane, some transcripts are also or even exclusively enriched near the ER surface (expansion shows genes with more than 2-fold enrichment at the ER). **B**, The ER localization of Oxa1 was determined with the split-GFP assay in wild type and Δ*djp1* cells that were grown to log phase in glucose medium. Oxa1 is trapped at the ER in Δ*djp1*. Mean values and standard deviations from *n*=3 independent biological replicates are shown. **C**, *HAC1* splicing in wild type cells grown to log phase in media with the indicated carbon sources was measured via RT-qPCR. *HAC1*^*i*^ levels were normalized to total *HAC1* levels. Mean values and standard deviations from *n*=5 independent biological replicates are shown. **D**, Wild type, Δ*ire1* and Δ*hac1* cells were grown to log phase in glucose, galactose and lactate media, washed and switched to glucose, galactose and lactate media in all combinations. Growth was monitored by OD_600_ measurement in a plate reader. Both UPR^ER^-deficient mutants showed impaired growth when the carbon source was switched to one that promotes higher levels of *HAC1* splicing in wild type cells. **E**, Wild type, Δ*ire1* and Δ*hac1* cells were plated on glycerol and 10 µl of a 3 M solution of the UPR^ER^-inducing agent dithiotreithol (DTT) were applied on a filter dish in the middle of the plate. Note the ring-like growth of the wild type around the filter dish. **F**, Wild type cells and cells that express *HAC1*^*i*^ from an estradiol-inducible *GAL* promoter were grown to log phase in glucose medium. They were washed, resuspended in either glucose or lactate medium supplemented with the indicated concentration of estradiol. Ectopic expression of low levels of *HAC1*^*i*^ result in better growth in lactate, but not in glucose medium. **G**, Schematic model for the role of the UPR^ER^ in mitochondrial protein biogenesis. A fraction of mitochondrial precursor proteins constantly localizes to the ER. Global changes in expression of mitochondrial genes increase the influx of precursors to the ER. Defects in protein import also elevate the levels of ER-resident mitochondrial precursors. In both cases, activation of the UPR^ER^ adjusts the proteostasis capacity of the ER.

We did not observe considerable fluorescence in our split-GFP assays without applying import stress. However, under steady state conditions, precursors might only very transiently localize to the ER because they can be efficiently rerouted to mitochondria with the help of the ER-resident J protein Djp1 in a process called ER-SURF ^15^. Loss of this pathway does not impair mitochondrial import *per se*, but would trap mitochondrial orphans at the ER. To test this, we employed our split-GFP assay in the ‘ER-trapping’ Δ*djp1* mutant and found accumulation of Oxa1 at the ER even under optimal growth conditions (Fig. 4B). Hence, there is indeed a constitutive flux of mitochondrial precursors to the ER in the absence of stress.

We therefore wondered whether the UPR^ER^ might be required to buffer fluctuations in the levels of ER-localized mitochondrial precursors under physiological conditions. Mitochondrial biogenesis is strongly dependent on the carbon source in the growth media: The levels and, hence, synthesis of many mitochondrial proteins are low on glucose, but considerably higher on raffinose, galactose, glycerol or lactate ^68-70^. In fact, we observed that the extent of steady state *HAC1* splicing was low when cells were grown on glucose, but elevated on all other carbon sources, particularly on galactose and glycerol (Fig. 4C). To assess the functional relevance of the UPR^ER^ under different states of mitochondrial metabolism, we grew wild type, Δ*hac1* and Δ*ire1* cells to exponential phase in liquid medium containing glucose, galactose or lactate as sole carbon source. Then we washed the cells, resuspended them in glucose, galactose and lactate medium in all possible combinations and monitored their growth (Fig. 4D). While there was no difference between wild type and ER-deficient strains when they remained in the media they were cultured in before, Δ*hac1* and Δ*ire1* mutants had problems to adapt when carbon sources were switched to a medium with higher levels of *HAC1* splicing. Likewise, ER-deficient strains grew well during exponential phase in glucose, but exhibited a phenotype at high optical densities, shortly before the cultures entered the stationary phase. At this point, yeast cells respond to the depletion of glucose and switch to respiratory metabolism, a growth phase called diauxic shift in which mitochondrial biogenesis is strongly induced ^71,72^. Hence, the UPR^ER^ is important when such a remodeling takes place.

Would a stronger UPR^ER^ help cells to adapt to respiratory growth conditions? Yeast cells grew better on respiratory media when exposed to moderate amounts of the reducing agent dithiothreitol (DTT), which is a known trigger of the UPR^ER^ and, consequently, is toxic for UPR-deficient strains (Fig. 4E). However, its beneficial effect on respiratory growth might – at least in part – also result from UPR^ER^-independent effects. We therefore sought to induce the UPR^ER^ directly without any stress treatment by expressing the spliced isoform of *HAC1* from a β-estradiol-inducible *GAL* promoter ^23^. Cells were precultured in glucose medium and, upon shift to either glucose or lactate medium, exposed to various β-estradiol concentrations, i.e. to different levels of *HAC1*^*i*^ expression. Indeed, cells grew better in lactate when *GAL-HAC1*^*i*^ was induced with up to 50 nM β-estradiol, while they were not affected when grown in glucose (Fig. 4F). Higher concentrations of β-estradiol delayed growth, consistent with earlier reports that overshooting UPR^ER^ activation can be toxic ^73,74^.

Obviously, a functional UPR^ER^ is not only important when mitochondrial protein import is blocked, but also maintains cellular fitness under physiological conditions with elevated mitochondrial biogenesis. We propose that a fraction of mitochondrial precursor proteins is always localizing to the ER, either transiently as part of the ER-SURF pathway or terminally mistargeted. When the influx of precursors is altered due to changes in gene expression or by mitochondrial dysfunction, the UPR^ER^ acts as a ‘rheostat’ and adjusts the protein folding and quality control components of the ER accordingly (Fig. 4G).

## Discussion

Precursor proteins that accumulate outside mitochondria impose a burden on cellular proteostasis. Many precursors remain in the cytosol ^31^ or end up in the nucleus ^75^, where chaperones and the proteasome mitigate the adverse effects of mistargeted proteins and eventually degrade them ^27,28,76,77^. Membrane proteins are particularly prone to misfolding and aggregation in an aqueous environment. Hence, their prolonged presence in the cytosol can be very hazardous for cells ^78,79^. We found in this study that cells can adsorb precursors of mitochondrial membrane proteins to the surface of the ER and employ the UPR^ER^ to buffer their elevated levels at the ER. Apparently, mitochondrial proteins associate with the ER even under physiological conditions. However, the accumulation of mitochondrial precursors at the ER is exacerbated by import defects as well as by metabolic stimuli that increase the expression of abundant mitochondrial enzymes, many of which are membrane proteins. Our observations identify the UPR^ER^ as an important cellular response to promote cellular fitness under such conditions, especially during the phase of adaptation.

There are numerous reasons why engaging the ER as a venue for buffering mitochondrial membrane proteins can be beneficial: (1) The large ER membrane provides a favorable environment for proteins with hydrophobic transmembrane domains that would otherwise misfold in the aqueous cytosol. (2) The ER has a remarkable capacity to prevent protein aggregation, even exceeding that of the cytosol for some classes of proteins ^80,81^. (3) Besides having chaperones that promote protein folding, the ER harbors an elaborate machinery for ER-associated protein degradation (ERAD). ER components have been found to participate in the degradation of cytosolic and, more recently, mitochondrial proteins ^82,83^. (4) ER and mitochondria share many components in their protein biogenesis and quality control systems, e.g. the Hsp40 co-chaperone Ydj1 or Cdc48/VCP/p97 and many of its cofactors ^36,84-86^. In addition, some organelle-specific factors of ER and mitochondria physically interact and functionally cooperate with each other ^87^. (5) Protein transfer between mitochondrial and ER membranes is possible via dedicated machineries that can extract mislocalized proteins from the membrane and set them back en route to the respective other organelle ^15,88-91^. (6) The close proximity of mitochondria and ER at membrane contact sites might facilitate the exchange of proteins between the two organelles. Interestingly, ER-mitochondria contact sites are enriched with ER chaperones and other UPR^ER^ effectors ^92^ and their loss activates the UPR^ER 93^. In addition, contact sites are crucial for the initiation of autophagy ^94,95^.

Based on the above considerations, it is possible that routing of mitochondrial precursors to the ER could be more than a mere ‘mistake’ in protein targeting, but rather an actively regulated quality control pathway. In line with this idea, our analyses show that SRP recognizes and binds nascent chains of some mitochondrial proteins, suggesting that a portion of the mitochondrial proteome is synthesized at the ER surface. Also the GET pathway (guided entry of tail-anchored proteins) was recently identified to be involved in ER targeting of mitochondrial tail-anchored proteins and some carrier proteins ^88,96^.

Our findings add to a significant body of observations linking the stress responses and homeostasis mechanisms of mitochondria and ER ^97^. Several processes connect mitochondrial and ER homeostasis in the context of stress: the flux of lipids between mitochondrial and ER membranes ^98^; the generation of ATP as well as reactive oxygen species by the respiratory chain ^99,100^; the transport of calcium ^16,101^; or the availability of building blocks for glycosylation of secretory proteins, provided by mitochondrial carbohydrate metabolism ^102^. We propose that mitochondria and ER are also linked in the management of mitochondrial biogenesis.

Our findings open the question which of the many components and pathways that are reinforced by the UPR^ER^ are the most important for the management of ER-localized mitochondrial proteins. It will be exciting to disentangle the exact contributions of storage and handling of misfolding-prone precursors, their transfer to mitochondria or their degradation at the ER surface.

We suggest that cells engage the ER and its proteostasis capacity – augmented by the UPR^ER^ when necessary – as a buffer for proteins that can’t be immediately imported into mitochondria. From there, they can be either degraded or kept on hold for a second attempt of mitochondrial import.

Therefore, we should consider to rethink the classical concept of ‘mislocalization’ as a problem that cells need to avoid. Rather, spatial sequestration (transient or terminal) of proteins to compartments other than those they are primarily targeted to might be a productive step in protein biogenesis. It will be exciting to explore this concept and the components that are involved in the future.

## Materials and Methods

### Yeast strains and plasmids

All yeast strains used in this study are listed in Supplementary Table S1 and were based on the wild type strain W303 or YPH499 ^103,104^. The *mia40-4* strain was a gift from Agnieszka Chacinska ^44^. Yeast strains were grown on YP medium (1% yeast extract, 2% peptone) or synthetic medium (0.17% yeast nitrogen base and 0.5% (NH_4_)_2_SO_4_) containing 2% glucose, 2% galactose, 2% raffinose, 2% glycerol or 2% lactate and supplemented with appropriate amounts of amino acids and nucleobases for selection.

pFA6a-ymNeongreen-*CaURA3* and pFA6a-ymScarletI-*CaURA3* were kindly provided by Bas Teusink (Addgene plasmids # 125703 and # 118457) ^105^. Genomic tagging with ymNeonGreen was performed by amplifying the ym-NeonGreen-CaURA3 cassette with overhangs homologous to the *OXA1* locus and transforming yeast cells using the lithium acetate/ss carrier DNA/PEG method ^106^. Genomic deletion of *IRE1* and *HAC1* in the *mia40-4* background was performed by amplifying a kanMX4 cassette from a pFA6a plasmid with overhangs homologous to the sequences up- and downstream of the genomic open reading frames of the target genes ^106^. Yeast cells were transformed with the PCR product and grown on plates containing 150 µg/ml G418 for selection. Deletions were confirmed by colony PCR on the targeted genomic loci.

The sequences of GFP^11^ (pSJ1321, pRS315-NOP1pr-GFP11-mCherry-PUS1) and GFP^1-10^ (pSJ2039, pRS316-NOP1pr-GFP1-10-SCS2TM) were a gift from Sue Jaspersen (Addgene plasmids # 86413 and # 86418) ^48^. Cloning of the split-GFP constructs into the pYX122, pYX142 and pNH605 plasmids used in this study was performed by Gibson Assembly with the HiFi DNA Assembly Master Mix (New England Biolabs, #E2621L) according to the manufacturer’s instructions. The GFP^11^ part was fused to the different proteins by integration of the sequence (5’ AGA GAT CAT ATG GTT TTG CAT GAA TAT GTT AAT GCT GCT GGT ATT ACT TAA 3’) into the corresponding primers. GFP^1-10^ was amplified from the plasmid (pSJ2039) with overhangs homologous to the end of the fusion partner and the plasmid pYX122.

### Isolation of RNA and RT-qPCR

RNA was extracted from yeast cells using either the acid phenol-chloroform method or an RNeasy Mini kit with on-column removal of DNA (Qiagen), both as previously described ^28^. In either case, 3 OD_600_×ml of cells were collected by centrifugation (17,000 × *g*, 3 min, 2°C), washed with prechilled water, snap-frozen in liquid nitrogen and stored at -80°C.

For acid phenol-chloroform extraction, cell lysates were prepared in lysis buffer (50 mM Tris/HCl (pH 7.0), 130 mM NaCl, 5 mM EDTA, 5% (w/v) SDS) with a FastPrep-24 5 G homogenizer (MP Biomedicals) with 3 cycles of 20 s, speed 6.0 m s^−1^, 120 s breaks, lysis matrix Y). RNA was purified with repeated extraction with acid phenol–chloroform (5:1, pH 4.5, two times) and 24:1 chloroform–isoamylalcohol (24:1). Then, 0.3 M sodium acetate (pH 5.5) was added, RNA was precipitated with ethanol and solubilized in water. DNA was removed using a Turbo DNA Free kit (Ambion) following the manufacturer’s instructions. RNA purity and concentration were assessed using a DeNovix DS-11 FX+ Fluorometer.

RT-qPCR was performed with a CFX96 Touch Real-Time PCR Detection System (Bio-Rad). 100 ng total RNA per 20 µl reaction were analyzed using the Luna Universal Probe One-Step RT-qPCR Kit (NEB, # E3006) in technical triplicates. cDNA was generated by reverse transcription for 10 min at 55°C. PCR amplification was then carried out under the following conditions: initial denaturation for 1 min at 95°C, followed by 45 cycles of 10 s at 95°C (denaturation) and 30 s at 60°C (extension). Primer-probe combinations for qPCR are listed in Supplementary Table 3. For the specific detection of the spliced isoform of *HAC1*, primers were chosen to flank the intron and the fluorescent probe spans the exon-exon junction (Fig. S1B). Primer efficiency was determined by measuring serial dilutions of pooled cDNA and only primer-probe combinations with an efficiency within 90% and 110% were used. *C*_*q*_ values were obtained with the Bio-Rad CFX Manager 3.1 with *C*_*q*_ Determination Mode set to “Single Threshold” and Baseline Setting set to “Baseline Subtracted Curve Fit”. Gene expression was normalized to the geometric mean of the expression values of the reference gene *TFC1* ^107^. Statistical significance was assessed with paired two-tailed Student’s *t*-test.

### Growth Assays

Growth curves were performed automated in a 96 well plate in technical triplicates using the ELx808 Absorbance Microplate Reader (BioTek). Precultures of 100 μl were inoculated at an OD_600_ of 0.1 in round bottom microtiter plates and sealed with an air-permeable membrane (Breathe-Easy; Sigma-Aldrich, St. Louis, MO). The growth curves started at OD_600_ 0.1 and incubated at 30°C for 72 h under constant shaking. The OD_600_ was measured every 10 min.

For the Halo assay, strains were grown in liquid YPD media to mid-log phase, washed and plated on YPG plates. A filter plate was placed onto the plate and soaked with 10 µl of a 3 M DTT solution. Plates were incubated at 30°C for 2 days.

### Preparation of Cell Extracts for Western Blotting

For whole cell lysates yeast strains were cultivated in selective lactate media. Clogger expression was induced by adding 0.5% galactose. After 4.5h, 2 OD_600_×ml of cells were harvested by centrifugation (5,000g, 5 min, RT) and washed with water. The cells were resuspended in 40 µl/OD_600_ 1x Laemmli buffer (125 mM Tris/HCl (pH 6.8), 5% SDS (w/v), 25% glycerol, 0.0005% bromophenol blue) and transferred to a screw-cap tube containing glass beads (0.5 mm). Cell were lysed using a FastPrep-24 5 G homogenizer (MP Biomedicals) with 3 cycles of 20 s, speed 6.0 m/s, 120 s breaks, lysis matrix Y. Cell extracts were boiled for 10 min at 96°C. Samples were stored at - 20°C until usage. An equal amount of lysate corresponding to 0.4 OD_600_×ml per sample was loaded on an SDS gel.

### Immunoblotting

Proteins were separated by size using discontinuous sodium dodecyl sulfate polyacrylamide gel electrophoresis (SDS-PAGE). They were transferred to a nitrocellulose membrane by semi-dry western blotting with blotting buffer (20 mM Tris, 150 mM glycine, 0.08% SDS (w/v), 20% methanol). To visualize the transferred proteins, the membrane was stained with Ponceau S solution (0.2% (w/v) Ponceau S, 3% (w/v) acetic acid) for 5 min. The membrane was cut in pieces to decorate against several antibodies at once and unspecific binding was blocked by incubation for 30 min in 5% milk in 1X TBS buffer (10 mM Tris/HCl (pH 7.5), 150 mM NaCl). The first antibodies were incubated over night at 4°C. The membrane was washed extensively with 1X TBS Buffer. Afterwards, the membrane was incubated for 90 min at room temperature with the secondary antibody containing the horseradish peroxidase (anti-Rabbit). The membrane was again washed extensively before ECL1 (100 mM Tris/HCl (pH 8.5), 0.044% (w/v) luminol, 0.0066% p-coumaric acid) and ECL2 (100 mM Tris/HCl (pH 8.5), 0.03% H_2_O_2_) solutions were mixed 1:1 and poured onto the membrane. Thereby chemo luminescence is produced by horseradish peroxidase coupled to the secondary antibody, which was detected on Super RX Medical X-Ray Films (Fuji) using the Optimax Type TR-developer.

### Antibodies

The antibodies for the use in immunoblotting of *S. cerevisiae* cell extracts were raised in rabbits using purified recombinant proteins. The secondary antibody was ordered from Biorad (Goat anti-Rabbit IgG (H+L)-HRP Conjugate #172-1019). Antibodies were diluted in 5% (w/v) nonfat dry milk-TBS (Roth T145.2) with the following dilutions: anti-Sod1 1:1,000, anti-Rip1 1:750, anti-Mdj1 1:125, anti-Rabbit 1:10,000. anti-Rip1 and anti-Mdj1 sera were a gift from Thomas Becker.

### Split-GFP Assay

Cells containing were transformed with one of the plasmids pYX142-Oxa1-GFP^11^, pYX142-Om45-GFP^11^, pYX142-Dld1-GFP^11^ or pYX142-Mia40-GFP^11^ in combination with either pYX122-Sec63-GFP^1-10^, pYX122-Oxa1-GFP^1-10^, pYX122-Mia40-GFP^1-10^ or pYX122-Ssa1-GFP^1-10^. All combinations contained also either the plasmid pYX233-*b*_*2*_-DHFR or the control plasmid pYX233-cyt DHFR. Cells were grown in selective medium containing 2% lactate to mid log phase. Mitoprotein-induced stress was induced by addition of 0.5% galactose for 4.5 h. 3 OD_600_×ml were harvested, resuspended in 100 µl medium containing 2% lactate, transferred into a black 96 well plate and centrifuged (5 min at 30 g). The fluorescence was measured with the excitation/emission wavelengths 485±15/530±20 nm in a fluorescence microplate reader (Clariostar, BMG labtech).

For the time course measurement of split-GFP fluorescence in a growing culture, the split-GFP cassette with Oxa1-GFP^11^ and Sec63-GFP^1-10^ was genomically integrated into the *LEU2* locus of yeast cells. In addition, a constitutively expressed *TEF1p*-ymScarletI was integrated into the *HIS3* locus. The cells were transformed with a pYX233 plasmid for either cytosolic DHFR or *b*_*2*_-DHFR expression and grown to mid-log phase in synthetic lactate medium. The cells were then diluted to an OD_600_ of 0.4 in 100 µl lactate medium with (inducing) or without (non-inducing) 0.5% galactose in a microtiter plate sealed with an air-permeable membrane (Breathe-Easy; Sigma-Aldrich, St. Louis, MO) in *n*=6 replicates. A WT strain not expressing no fluorescent protein was used for correction of the background fluorescence of cells or media. The plate was incubated at 30°C under recurrent shaking in a ClarioStar spectrofluorometer (BMG Labtech) and fluorescence was measured every 10 min with the following excitation/emission wavelengths: 485±15/530±20 nm for split-GFP and 580±15/631±36 nm for ymScarletI. Background fluorescence was substracted and the split-GFP signal was divided by the ymScarletI signal to control for growth and overall translation. The average fluorescence intensity at timepoint 0 was set to 1.

### Fluorescence microcopy

To analyze the localization of the fluorescence signal in the different split-GFP combinations, mid log phase cultures were shifted to media containing 0,5% galactose to induce the expression of the *b*_*2*_-DHFR clogger or cytosolic DHFR as control. After centrifugation of 1 OD_600_×ml of cells (1 min at 16.000 g at RT), the cells were resuspended in 50 µl sterile water. The cell suspension was transferred to a microscope slide for fluorescence imaging using the HCX PL APO 63x oil immersion objective of a Leica TCS SP5II confocal laser scanning microscope. GFP was excited at 488 nm and emission was detected by a photomultiplier through a 530/30-nm band pass filter. Microscopy images were processed using Leica software LAS X (v3.3) and Fiji (v2.1.0).

### Ribosome Profiling

#### Library Preparation

Yeast cultures were grown to mid-log phase in minimal medium containing 2% lactate. Expression of *b*_*2*_-DHFR or cytosolic DHFR was induced by addition of 0.5% galactose for 4.5 h. Cells were harvested by vacuum filtration (pore size 0.45 µm). In one out of three independent replicates, 100 µg/ml cycloheximide was added to the yeast culture 2 min before harvesting and lysis to inhibit translation elongation, while in the other replicate, cells were not in contact with cycloheximide prior to cell lysis. Cells were flash-frozen in liquid nitrogen and lysed in a mixer mill (Retsch, MM 301) in lysis buffer (20 mM Tris/HCl (pH 7.4), 140 mM KCl, 1.5 mM MgCl_2_, 0.5 mM DTT, 100 µg/ml cycloheximide, 1% (v/v) Triton X-100) in 50 ml stainless steel grinding chambers under cryogenic conditions for 1 min at 20 Hz. Lysates were thawed in a water bath at room temperature, immediately followed by centrifugation at 15,000 g at 4°C for 10 min. RNA concentration was quantified with a NanoDrop fluorometer (absorbance at 260 nm) and RNA digestion was performed with RNase I (Ambion, #AM2294, 2.5 µl / mg RNA) for 45 min at room temperature. Digestion was stopped by the addition of SUPERase·In RNase inhibitor (Ambion, #AM2696, 2 µl / 100 µl digestion). Ribosomes were isolate by centrifugation through a 25% (w/v) sucrose cushion in a

TLA 100.2 rotor (Beckman) at 72,000 rpm for 20 min at 4°C. RNA was extracted from the ribosomal pellet using the hot SDS-Phenol-Chloroform method and 24-35 nt ribosome footprints were size selected on a 15% (w/v) polyacrylamide TBE-urea gel. Ribosomal RNA was removed with the RiboZero Gold kit (Illumina). Sequencing libraries were then prepared as previously described ^108^. Libraries were quantified by qPCR (Kapa Biosystems) and sequenced using a HiSeq 4000 (Illumina).

#### Data analysis

Sequencing reads were demultiplexed with Illumina CASAVA v1.8 and adaptor sequences were trimmed using Cutadapt v2.8. Reads that mapped to ribosomal RNAs were removed using Bowtie v.1.2.3 ^109^ and remaining reads were aligned to the yeast reference genome obtained from the *Saccharomyces* genome database (genome release R64-2-1).

For each read, reads were summed at each nucleotide by customized python scripts. Metagene analysis was performed separately on each fragment length to remove lengths that did not exhibit the 3-nucleotide periodicity that is characteristic for ribosome footprints. Each of the remaining reads was assigned to the first A-site nucleotide. To this end, a nucleotide offset from the 5’ end of each fragment length was empirically determined, using the characteristic high ribosome density at the start codon. Nucleotide reads at each codon were then summed and used for all downstream analysis.

Gene-level differential expression analysis was performed using HTSeq ^110^ and the DESeq2 package ^111^ within the Bioconductor v3.12 project in the statistical programming language R v.4.0.3 ^112^.

### Analysis of published datasets on mRNA localization

The dataset on translation close to the ER and mitochondrial surface in yeast was obtained from Jan et al. (2014)^59^. In this study, the authors fused the biotin ligase BirA to Sec63 (ER) or Om45 (mitochondrial) and pulled down ribosomes that were biotinylated after a short pulse of biotin and translation inhibition with cycloheximide (CHX). The genes were filtered for those that code for mitochondrial proteins according to ^70^ and log_2_ fold enrichments of ribosome footprints at the ER (7 min CHX) or mitochondrial membranes (2 min CHX) over total ribosome footprints were plotted.

The dataset on transcript localization in human cells was obtained from Fazal et al. (2019)^61^. Here, the authors used the biotin ligase APEX2 fused to proteins of different cellular localizations to directly biotinylate RNA. Mitochondrial genes were filtered according to MitoCarta 3.0 ^113^ and log_2_ fold enrichment of ER- or mitochondria-localized transcripts over total transcripts were plotted.

The dataset on the SRP-bound translatome in yeast was obtained from Chartron et al. (2016)^63^. The authors compared ribosome-nascent chain complexes purified by pulldown of SRP to total ribosomes by ribosome profiling. Genes coding for secretory, cytosolic and mitochondrial proteins were filtered according to the author’s categorization and the distribution of the log_2_ fold enrichment of SRP-bound polysomes over total ribosome footprints was plotted.

### Data and material availability

The data produced in this study are presented in this published article and its supplementary material. The ribosome profiling data on clogger-expressing yeast cells are deposited into GEO ^114^ with accession number GSE172017.

All yeast strains, plasmids and primers used in this study are listed in Supplementary Tables 1-3 and are available from the authors upon request. The plasmids pYX233 DHFR and pYX233 *b*_*2*_-DHFR for expression of the mitochondrial clogger are available via Addgene (plasmids #163761 and #163759).

## Author contributions

F.B. and J.F. conceived and supervised the study. F.B. and K.C.S. prepared the ribosome profiling libraries and performed the bioinformatics analysis of the sequencing data. K.K., L.K., C.G. and F.B. generated constructs and strains. K.K., L.K. and C.G. performed *in vivo* experiments. K.K. and F.B. analyzed *HAC1* splicing by RT-PCR. K.K. established and performed the split-GFP assay. K.K. performed fluorescence microscopy and K.K. and K.G.H. analyzed the results. K.G.H. and F.B. analyzed ribosome profiling data on localized and SRP-bound translation of mitochondrial proteins. K.K., C.G., L.K., K.G.H., J.M.H., J.F. and F.B. analyzed the data. F.B. wrote the manuscript with input from all authors.

## Acknowledgements

Sequencing was performed at the UCSF Center for Advanced Technology. We thank Sabine Knaus, Andrea Trinkaus and Natalia M. Barbosa for assistance with the experiments and Torsten Möhlmann for help with the microscopy. We thank T. Kelly Rainbolt and Sebastian Schuck for discussions and comments on the manuscript. This project was funded by grants from the Landesforschungsinitiative Rheinland-Pfalz BioComp (to F.B. and J.M.H.), the Deutsche Forschungsgemeinschaft (DE2803/10 to J.M.H.), the US National Institutes of Health (NIH GM56433 to J.F. and AG047126 to K.C.S.), the Glenn Foundation for Medical Research (Postdoctoral Fellowship to K.C.S.), the Company of Biologists (Travelling Fellowship JCSTF181148 to F.B.), the European Molecular Biology Organization (Long-Term Fellowship ALT 762-2019 to K.G.H.), the Helen Hay Whitney Foundation (F-1240 to K.G.H.) and the Joachim Herz Stiftung (to F.B. and C.G.).

## Supplementary Figures

**Supplementary Figure 1.**
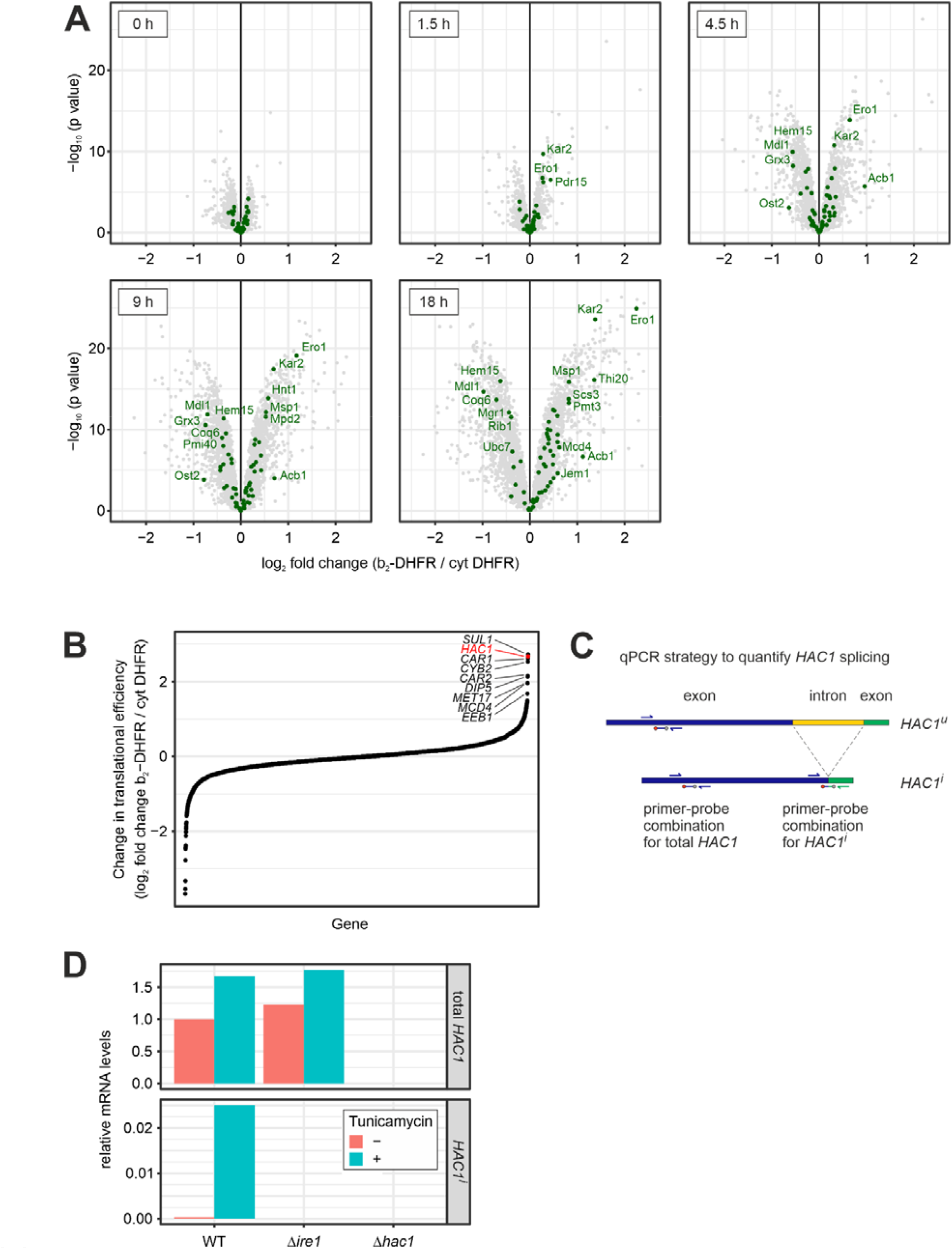
Detection of UPR^ER^ induction with mass spectrometry and RT-qPCR. **A**, Protein levels in clogger-expressing versus control cells after different times of induction were measured by quantitative mass spectrometry ^28^. Highlighted are proteins which are reported targets of the UPR^ER 21^. Data from *n*=3 independent biological replicates are shown. The data for 18 h are the same as shown in Fig. 1E. **B**, The change in translational efficiency after 4.5 h clogger expression was calculated for all genes measured in both the RNA-seq ^28^ and Ribo-Seq on clogger-expressing cells by dividing the translatome fold change by the transcriptome fold change. **C**, Schematic depiction of the primer-probe combinations used to quantify total *HAC1* as well as spliced *HAC1*^*i*^ mRNA levels via RT-qPCR. **D**, Wild type, Δ*ire1* and Δ*hac1* cells were grown in presence or absence of 1 µg/ml tunicamycin and *HAC1* and *HAC1*^*i*^ levels were analyzed with the primer-probe assay shown in C. As expected, *HAC1*^*i*^ levels increased in wild type cells treated with tunicamyin, but no *HAC1*^*i*^ was detected in cells lacking *HAC1* or *IRE1*, confirming the specificity of the RT-qPCR assay.

**Supplementary Figure 2.**
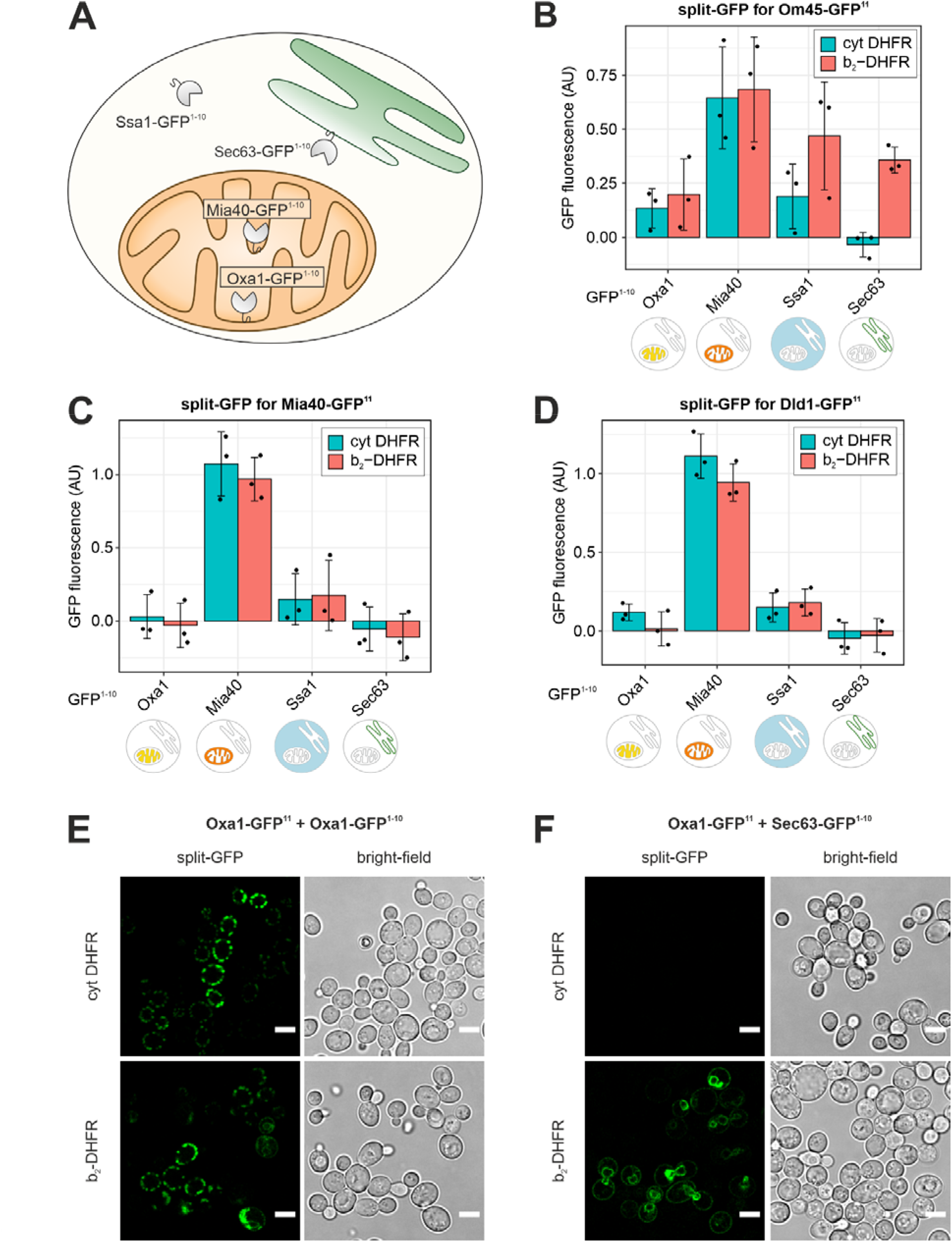
A split-GFP assay to assess the subcellular localization of mitochondrial precursor proteins. **A**, The GFP^11^ fragment was fused to Oxa1, Om45, Mia40 and Dld1, and the GFP^1-10^ reporter was fused to Oxa1 (mitochondrial inner membrane, matrix side), Mia40 (mitochondrial inner membrane, IMS side), Sec63 (ER membrane, cytosolic side) and Ssa1 (cytosol). **B-D**, The split-GFP constructs described in A were co-expressed with *b*_*2*_-DHFR or cytosolic DHFR and fluorescence was measured with a Clariostar plate reader. Under non-stressed conditions (expression of cytosolic DHFR), the split-GFP signals recapitulated the known localizations of Oxa1, Om45, Mia40 and Dld1. Under mitoprotein-induced stress (*b*_*2*_-DHFR expression), Om45-GFP^11^ also evoked a fluorescence signal when combined with Sec63-GFP^1-10^ and Ssa1-GFP^1-10^, indicating accumulation at the cytosolic side of the ER membrane. **E**, Fluorescence microscopy of cells expressing Oxa1-GFP^11^ and Oxa1-GFP^1-10^ and either *b*_*2*_-DHFR or cytosolic DHFR after 4.5 h of induction. Scale bar, 5 µm. **F**, Fluorescence microscopy of cells expressing Oxa1-GFP^11^ and Sec63-GFP^1-10^ and either *b*_*2*_-DHFR or cytosolic DHFR after 4.5 h of induction. Scale bar, 5 µm. GFP Images are identical to those in Figure 3E.

**Supplementary Figure 3.**
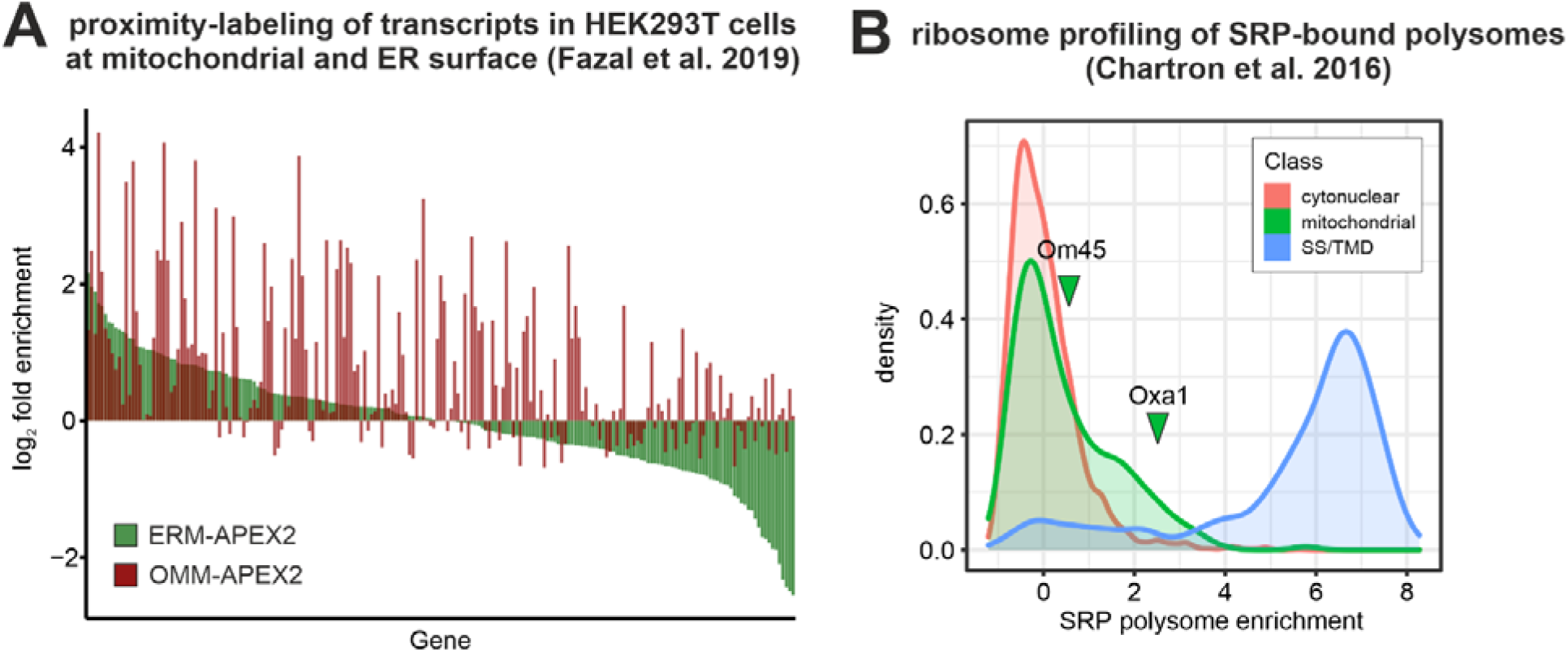
Certain mitochondrial proteins are synthesized close to the ER surface and recognized by SRP. **A**, Data from Fazal et al. (2019)^61^ on the subcellular distribution of mRNA in HEK293T cells. The biotin ligase APEX2 was localized to the ER or mitochondria and biotinylated mRNAs were purified and sequenced (APEX-Seq). For all mitochondrial proteins in the dataset, the log_2_ enrichment of mRNAs at the ER membrane (ERM-APEX2) and the mitochondrial outer membrane (OMM-APEX2) over the total mRNAs are shown. While most mRNAs localize to the mitochondrial membrane, some transcripts are also enriched near the ER surface. **B**, Data from Chartron et al. (2016)^63^ on the SRP-bound translatome in yeast. SRP was immune-purified from cell lysates and the co-isolated ribosome-nascent chains complexes were analyzed by ribosome profiling. The distribution of the log_2_ fold enrichment SRP-bound ribosome-nascent chain complexes over total ribosomes is shown for cytonuclear and mitochondrial proteins and proteins that carry a signal sequence or transmembrane domain for ER targeting (SS/TMD). Some mitochondrial proteins, including Oxa1, are bound by SRP.

## Supplementary Tables

**Supplementary Table 1**. Yeast strains used in this study.

**Supplementary Table 2**. Plasmids used in this study.

**Supplementary Table 3**. Primers used in this study.

